# Discriminating betacoronavirus receptor usage across subgenera using protein structure prediction and molecular dynamics

**DOI:** 10.64898/2026.09.25.754509

**Authors:** Shruthi S. Garimella, Guadalupe Lauro, Vardhan Peddamallu, Parth R. Bandivadekar, Anugraha Thyagatur, Roland Faller, Surl-Hee Ahn, Priya S. Shah

## Abstract

A critical step in the emergence of a virus is the ability of the viral protein to bind a host receptor and mediate cell entry. For many coronaviruses, this interaction occurs between the Spike S1 subunit and the human ACE2 receptor. Whether this binding interface can be computationally distinguished across unstudied viruses without experimentally resolved protein structures remains an open question. We predicted how 28 emerging coronaviruses may bind to human ACE2 using structural predictions, static interaction prediction programs, and molecular dynamics simulations. To screen the emerging coronaviruses, we predicted a library of S1 structures using AlphaFold. These predicted structures were then used to model the S1-ACE2 interaction with AlphaFold, ClusPro, and HADDOCK. We used known ACE2-binding sarbecoviruses as positive controls and coronaviruses that bind other receptors as negative controls to threshold predicted binding. Contact analysis quantified the predicted binding and revealed that these static interaction prediction methods varied in discriminative power. Less restrained static predictions separated binders from non-binders, whereas heavily restrained docking did not, potentially forcing an interaction where none should exist. This analysis highlighted an emerging coronavirus, Zhejiang2013, as a potential ACE2 binder. We used molecular dynamics simulations to further assess the static predictions and model the interaction over time. Overall, our results indicate that Zhejiang2013 exhibits dynamic interaction patterns consistent with ACE2 binding. Given that two ACE2-binding coronaviruses have caused global pandemics within the past two decades, identifying potential ACE2 binders is critical for early warning and pandemic preparedness.

## Introduction

Nearly every major pandemic of the past century has been caused by RNA viruses ^1^. A pandemic begins when a pathogen escapes its point of origin and spreads from a single geographical region to whole countries and continents simultaneously, disrupting lives and livelihoods. Often, these outbreaks start with a zoonotic spillover followed by amplification through human transmission and eventual widespread dissemination ^2^. As the world continues to urbanize and human populations encroach further into historically undisturbed areas, our increased interactions with wildlife increase the chances of new spillover events. Public health responses have focused on reacting to outbreaks through containment, vaccination, and therapeutic interventions. Anticipating spillover threats before they emerge requires understanding how a virus begins to infect a new host.

Coronaviruses are a clear example of why this matters. In the last 20 years, three major outbreaks have been caused by coronaviruses ^3^. Severe acute respiratory syndrome coronavirus (SARS-CoV, now referred to as SARS-CoV-1) was first identified in southern China in November 2002, marking the emergence of the first pandemic of the 21^st^ century ^4–6^. Middle East respiratory syndrome coronavirus (MERS-CoV) emerged in 2012 ^7^. Most recently, in December 2019, SARS-CoV-2 emerged to cause the COVID-19 pandemic, one of the deadliest in recent history ^8–11^. These events have firmly established coronaviruses as a persistent and pressing threat, underscoring the importance of assessing their potential for future emergence.

Coronaviruses are positive-sense, single-stranded RNA viruses with one of the largest viral RNA genomes, at around 30 kilobases. Such a large genome is replicated by an RNA-dependent RNA polymerase with an estimated error rate of around 1 mutation per 10,000 nucleotides. An exoribonuclease is also encoded to maintain genome fidelity ^12–14^. The frequent template switching during replication increases the likelihood of recombination events ^15^. This drives rapid genetic diversity, increasing the likelihood of virus emergence. For instance, recombination among different Spike sequences can lead to the emergence of bat sarbecoviruses ^12,16,17^. This, coupled with the pandemic potential of coronavirus spillover, has warranted research into the key interactions necessary for spillover ^2,4,18,19^.

There are only seven known coronaviruses that can infect humans, all belonging to either the alpha- or betacoronavirus genera. Human coronaviruses (HCoVs) 299E and NL63 from the alphacoronavirus genus cause the “common cold” ^20–23^. The betacoronavirus genus has five subgenera, three of which contain human-infecting coronaviruses: embecovirus, merbecovirus, and sarbecovirus ^15^. Merbecoviruses include MERS-CoV, and sarbecoviruses include SARS-CoV-1 and SARS-CoV-2, which have caused respiratory virus outbreaks and pandemics ^1,3,24^. Embecoviruses include HCoV-OC43 and HCoV-HKU1, which cause the “common cold” ^20,25–27^.

The interaction between Spike and a human receptor is a primary gatekeeping event for coronavirus emergence. ACE2 usage is common and an ancestral trait among sarbecoviruses, though there are several clades within the sarbecovirus subgenus that do not use ACE2 ^4,28–31^. Within the Spike protein, the receptor binding domain (RBD) in the S1 subunit is responsible for binding ACE2 ^32,33^. Some sarbecovirus RBDs require only a few mutations to bind human ACE2 ^12,30,34–36^. Binding to human ACE2 has been studied in detail for SARS-CoV-2 mutants, elucidating key residues and structural characteristics critical for binding ^35,36^. Studying emerging coronaviruses for their potential to bind human ACE2 can help us assess the risk they pose for emergence.

Computationally predicting protein binding between coronaviruses and potential receptors can expedite the screening and, consequently, prioritize viral proteins for downstream experimental testing. Advances in deep-learning based protein prediction tools and more accessible web-based servers, such as AlphaFold Server, ClusPro, and HADDOCK, provide opportunities to study virus-receptor protein interactions even when experimental data are limited. The AlphaFold Server, based on AlphaFold3, predicts the structure of a protein complex from the amino acid sequences of separate chains ^37,38^. Hereafter, we refer to this approach in AlphaFold3 to predict the protein-protein complex as Multimer. Multimer simply requires the amino acid sequences, which can be helpful for predicting proteins and complexes when little structural information is available. ClusPro applies rigid-body docking using Fast Fourier Transform to evaluate thousands of orientations of the two proteins and scores each orientation using van der Waals forces, electrostatics, and hydrophobicity ^39–43^. It filters out the lowest energy complexes until it determines the most physically plausible orientation. HADDOCK further integrates known information through restraints or binding interfaces to guide more biologically informed docking, serving as a semi-flexible docking platform ^44,45^. These prediction tools offer an initial view of potential binding orientations; however, these static predictions fail to capture the intrinsic flexibility, intermolecular forces, and dynamics of proteins on any time scale. Molecular dynamics (MD) simulations can validate static predictions by allowing complexes to evolve over time under physiological conditions. MD helps assess the stability and plausibility of static complexes, providing a more realistic understanding of protein-protein interactions and the intermolecular forces involved.

In this study, we aimed to determine how well protein structure and static interaction prediction methods can discriminate ACE2 receptor engagement across various betacoronavirus S1 proteins. We achieved this by compiling a representative set of betacoronaviruses across three subgenera, with some predicted and known to bind ACE2, and some that do not, and applying a computational pipeline to these representative sequences. We compared Multimer, ClusPro, and HADDOCK and found that ClusPro best recapitulated known ACE2 binding predictions. We also identified a hibecovirus, Zhejiang2013, as a previously unreported candidate ACE2 binder. We then prioritized this candidate for MD simulations to assess the physical feasibility of the binding predictions. Through this systematic approach, we determined that Zhejiang2013 has potential to bind human ACE2 similarly to known pandemic-causing coronaviruses. Overall, this work demonstrates that accessible computational tools can screen and prioritize emerging antigens for downstream testing as we shift from reactive to proactive surveillance of viral threats.

## Results

### Selection of coronaviruses to study

Our overall goal was to predict which emerging betacoronavirus S1s bind to human ACE2 using protein structure prediction, static protein docking, and sophisticated MD simulations (Figure 1A). We focused on the S1 region to balance the difficulties associated with modeling full-length Spike across different lineages while still incorporating nuances that may not be captured with RBD alone, since its location may vary across viruses ^32,46^. Previous work identified a set of sarbecoviruses and merbecoviruses as emerging coronaviruses, with varying predictions of receptor usage ^31^. Using this previous work as a reference, we compiled the S1 and RBD sequences for 28 coronaviruses from NCBI (Table 1). Our selections were made to test known ACE2 binders (positive controls), known ACE2 non-binders (negative controls), and a sampling of coronaviruses for which ACE2 binding/non-binding had been predicted based on phylogeny, but not tested experimentally (Figure 1B-C). We also included a hibecovirus, Zhejiang2013, as a distantly related coronavirus for which no ACE2 binding predictions or studies have been performed.

**Figure 1.**
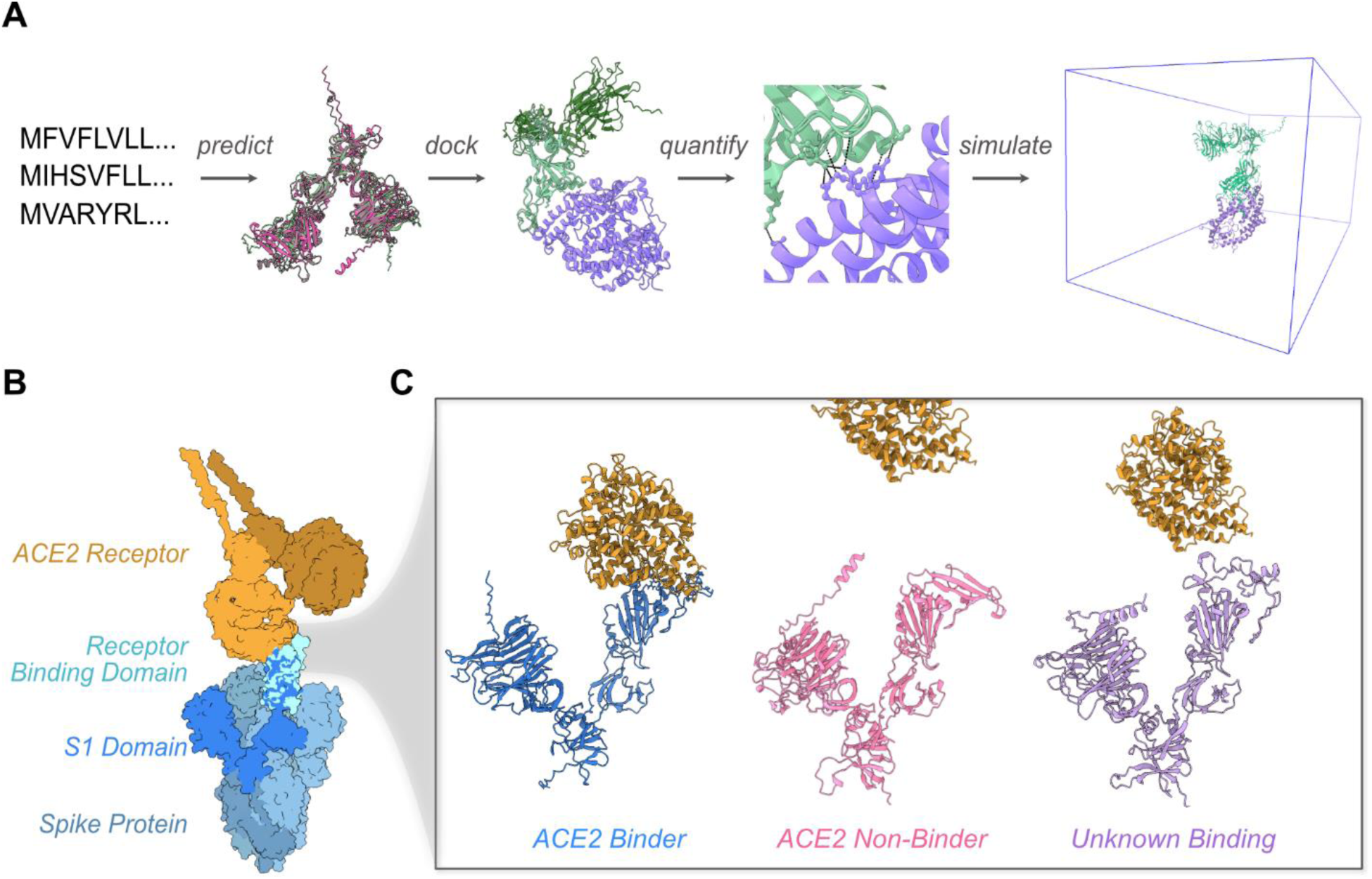
Overview of the study. A) Workflow of protein and interaction prediction to determine receptor binding. B) Estimated Spike-ACE2 binding using SARS-CoV-2 Spike trimer and human ACE2 dimer using PDB structures 6VXX, 61MD, 6M0J, and 6VSB. C) Example of coronavirus S1s as ACE2 binders, non-binders, and unknowns tested in this pipeline. AlphaFold predictions, from left to right, of S1 domains from sarbecovirus binder (SARS-CoV-2), merbecovirus non-binder (MERS-CoV), and hibecovirus with unknown binding (Zhejiang2013) subgenera.

**Table 1.** Compilation of coronaviruses used to test human ACE2 binding. We computationally tested 24 sarbecoviruses, three merbecoviruses, and one hibecovirus. For each virus, we use a shorthand abbreviation hereafter. The NCBI accession number is listed for each Spike protein, along with whether it binds human ACE2. Pandemic- and outbreak-causing coronaviruses that bind ACE2 are marked with *, and non-binders are noted with **.

| Subgenus | Virus Name | Abbreviation | NCBI Accession | ACE2 binder<br>30,31,34,35,47-49 |
| --- | --- | --- | --- | --- |
| Sarbecovirus | SARS-CoV-1 | SARS-CoV-1 | YP_009825051.1 | Yes* |
|  | SARS-CoV-2 | SARS-CoV-2 | QHD43416.1 | Yes* |
|  | LYRa11 | LYRa11 | KF569996 | Yes |
|  | LYRa3 | LYRa3 | KF569997.1 | Yes |
|  | RaTG13 | RaTG13 | QHR63300.2 | Yes |
|  | BetaCoV/P2V | P2V | QIQ54048.1 | Yes |
|  | Bat SARS-like coronavirus Rs3367 | Rs3367 | AGZ48818.1 | Yes |
|  | Bast SARS-like coronavirus Rs7327 | Rs7327 | ATO98231.1 | Yes |
|  | Bat coronavirus BtRs BetaCoV/YN2018B | YN2018B | QDF43825.1 | Yes |
|  | Bat SARS coronavirus Rm1/2004 | Rm1 | ABD75332.1 | Yes |
|  | Bat SARS coronavirus Rs672/2006 | Rs672 | ACU31032.1 | Yes |
|  | Bat SARS coronavirus HKU3 12 | HKU3 | ADE34812.1 | Yes |
|  | Bat coronavirus BtRf BetaCoV/SX2013 | SX2013C | AIA62300.1 | No |
|  | Bat SARS-like coronavirus YNLF31C | YNLF31C | AKZ19076.1 | No |
|  | Bat SARS-like coronavirus YNLF34C | YNLF34C | AKZ19087.1 | No |
|  | Bat coronavirus JTMC15 | JTMC15 | ANA96027.1 | No |
|  | SARS related coronavirus BtKY72 | BtKY72 | APO40579.1 | Yes |
|  | Bat SARS-like coronavirus Rs4255 | Rs4255 | ATO98193.1 | Yes |
|  | Bat coronavirus BtR1 BetaCoV/SC2018 | SC2018 | QDF43815.1 | No |
|  | Bat coronavirus BtRs BetaCoV/YN2018A | YN2018A | QDF43820.1 | Yes |
|  | Bat coronavirus BtRs BetaCoV/YN2018C | YN2018C | QDF43830.1 | Yes |
|  | Bat coronavirus BtRs BetaCoV/YN2018D | YN2018D | QDF43835.1 | Yes |
|  | Pangolin coronavirus Pang17 | Pang17 | QIA48632.1 | Yes |
|  | Bat SARS-like coronavirus W1V1 | W1V1 | AGZ48831.1 | Yes |
| **Merbecovirus** | MERS-CoV | MERS-CoV | NC_019843.3 | No** |
|  | Tylonycteris bat coronavirus HKU4 | HKU4 | YP_001039953.1 | No |
|  | Bat coronavirus BtCoV133/2005 | BtCoV133 | ABG47052.1 | No |
| **Hibecovirus** | Bat Hp betacoronavirus/Zhejiang2013 | Zhejiang2013 | AIL94216.1 | Unknown |

### Sequence alignment

We first compared the primary protein sequences of these S1 and RBD sequences using multiple sequence alignment, pairwise sequence identity calculations, and hierarchal clustering. We annotated which sequences were predicted/known to be ACE2 binders or non-binders (Figure 2). ACE2 non-binding sarbecoviruses formed one subgroup. Sarbecovirus ACE2 binders formed another subgroup. The ACE2 non-binding hibecovirus and merbecoviruses formed an outgroup.

**Figure 2.**
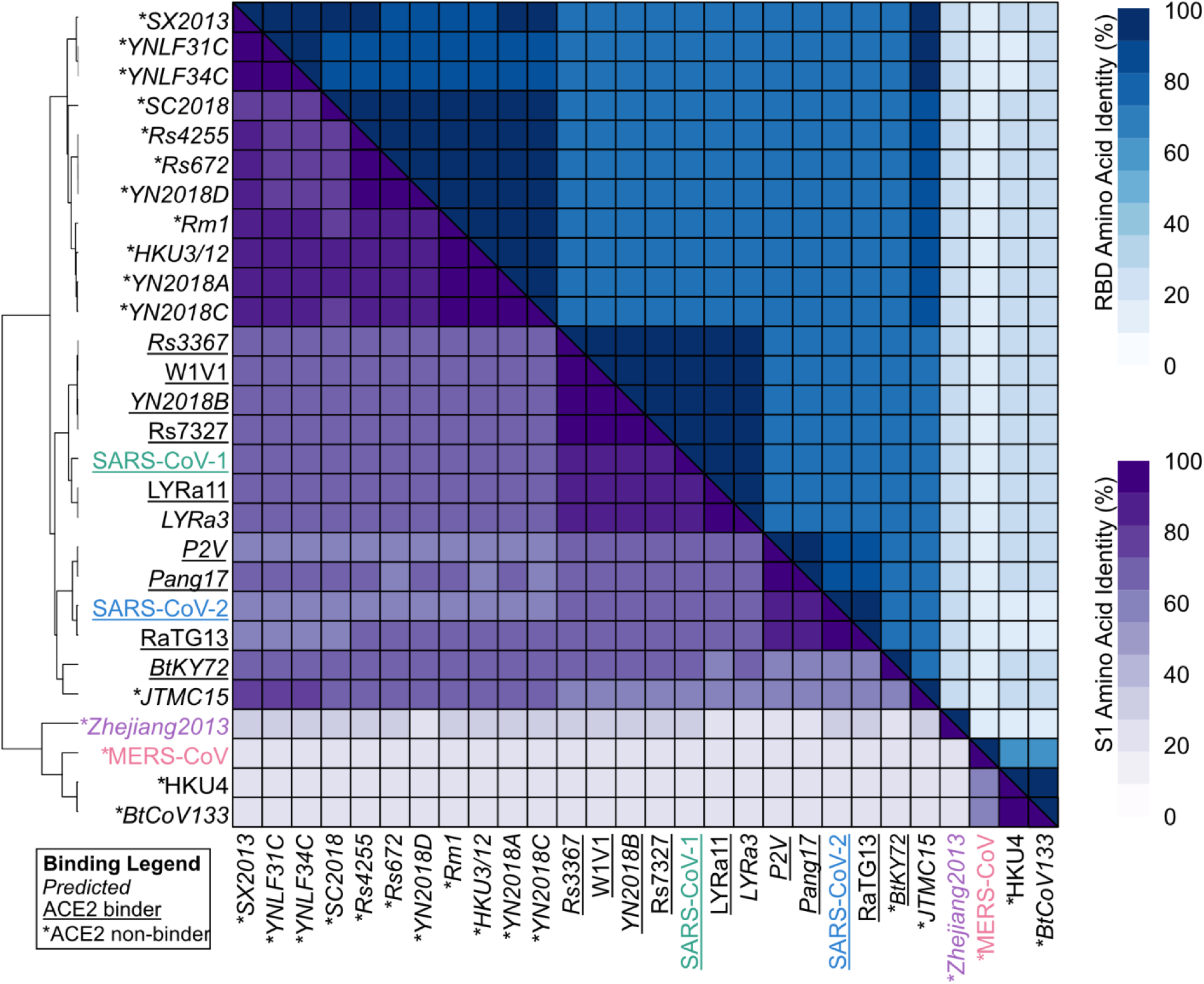
Sequence alignment heatmap of S1 (purple) and RBD (blue) using pairwise alignment of each betacoronavirus.

Overall, we observed at least 60% sequence identity in the S1 across all sarbecoviruses (Figure 2). Merbecoviruses and the hibecovirus showed much lower similarity to sarbecoviruses and to each other. The merbecoviruses only had a 20% identity to either subgenus. The hibecovirus shared about 27% sequence identity to sarbecoviruses that bind ACE2, 27% identity to sarbecoviruses that do not bind ACE2, and 20% identity to merbecoviruses that do not bind ACE2 (Table S1). The RBD lies near the middle of the S1 domain (Figure 1B). This position remained relatively consistent across the set of viruses. Trends observed with the S1 region were largely recapitulated by RBD sequence alignments (Figure 2 and Table S2).

### Protein structure prediction

We next predicted protein structures using AlphaFold (Figure 1A) ^50^. Visual inspection of the predicted proteins showed that the S1 structures had a relatively conserved shape (Figure 3). Merbecoviruses generally contained an additional beta sheet in their RBD compared to the sarbecoviruses and the hibecovirus. Additionally, the hibecovirus contained more intrinsically disordered regions in the RBD.

**Figure 3.**
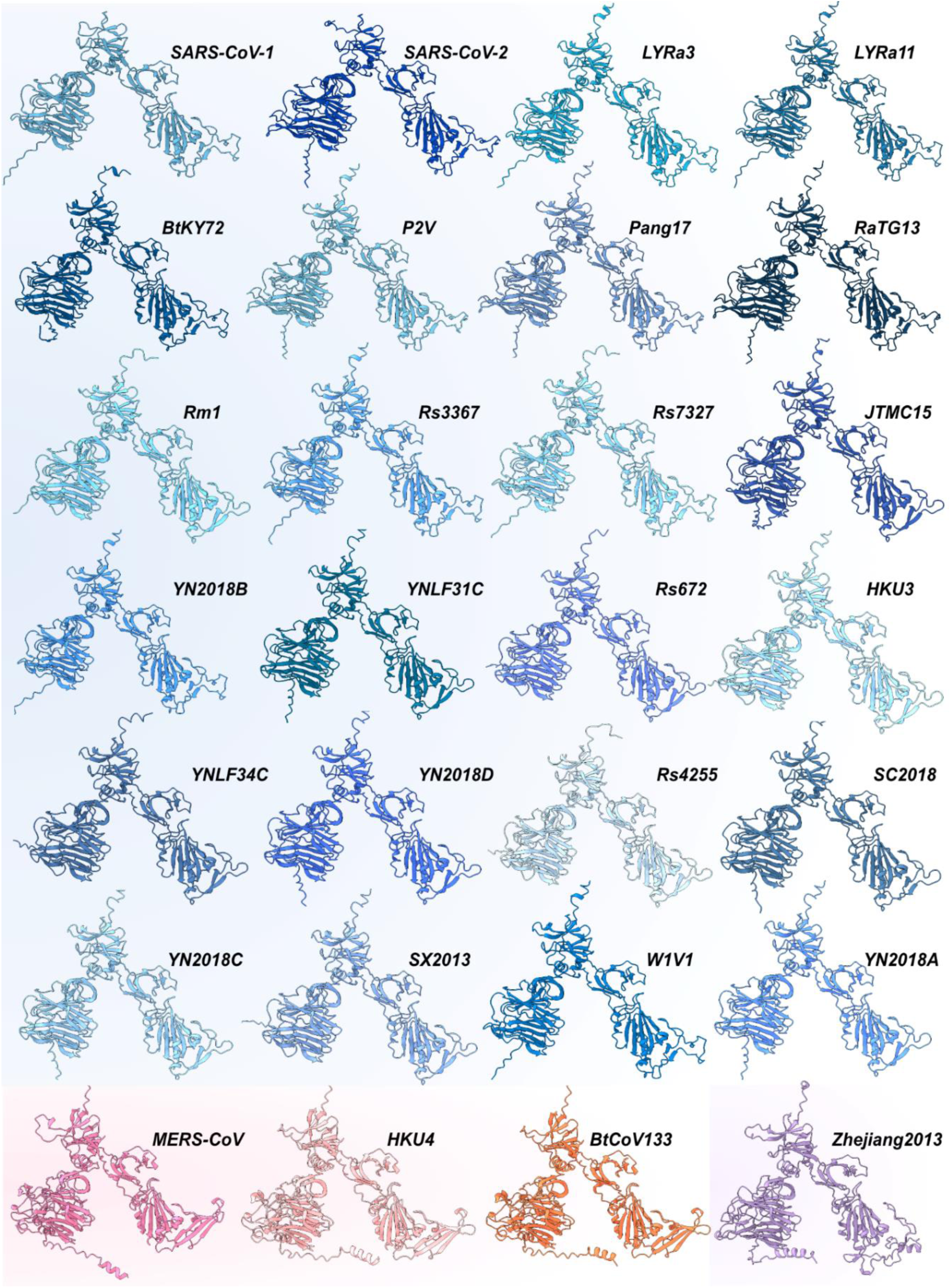
AlphaFold protein predictions of betacoronavirus S1 structures. Structures are oriented so that the RBD is towards the bottom right of the S1 domain. From left to right, the first 24 structures in blue are sarbecoviruses, the next three structures in pink are merbecoviruses, and the last structure in purple is a hibecovirus.

Using these structures, we examined the structural similarity between the proteins. We used US-align to obtain a template modeling (TM)-score to assess whether the trends observed with the sequence alignment are recapitulated with the protein structures. The TM-score is a length-independent metric that focuses on global alignment rather than local structural differences. A score of 1 indicates that two proteins are a perfect match, whereas anything above 0.5 can indicate similarly folded proteins ^51,52^. The trends observed for sequence identity (Figure 2) were also evident with structure similarity (Figure 4). As expected, sarbecoviruses shared more similarity with each other than with the hibecovirus and merbecoviruses.

**Figure 4.**
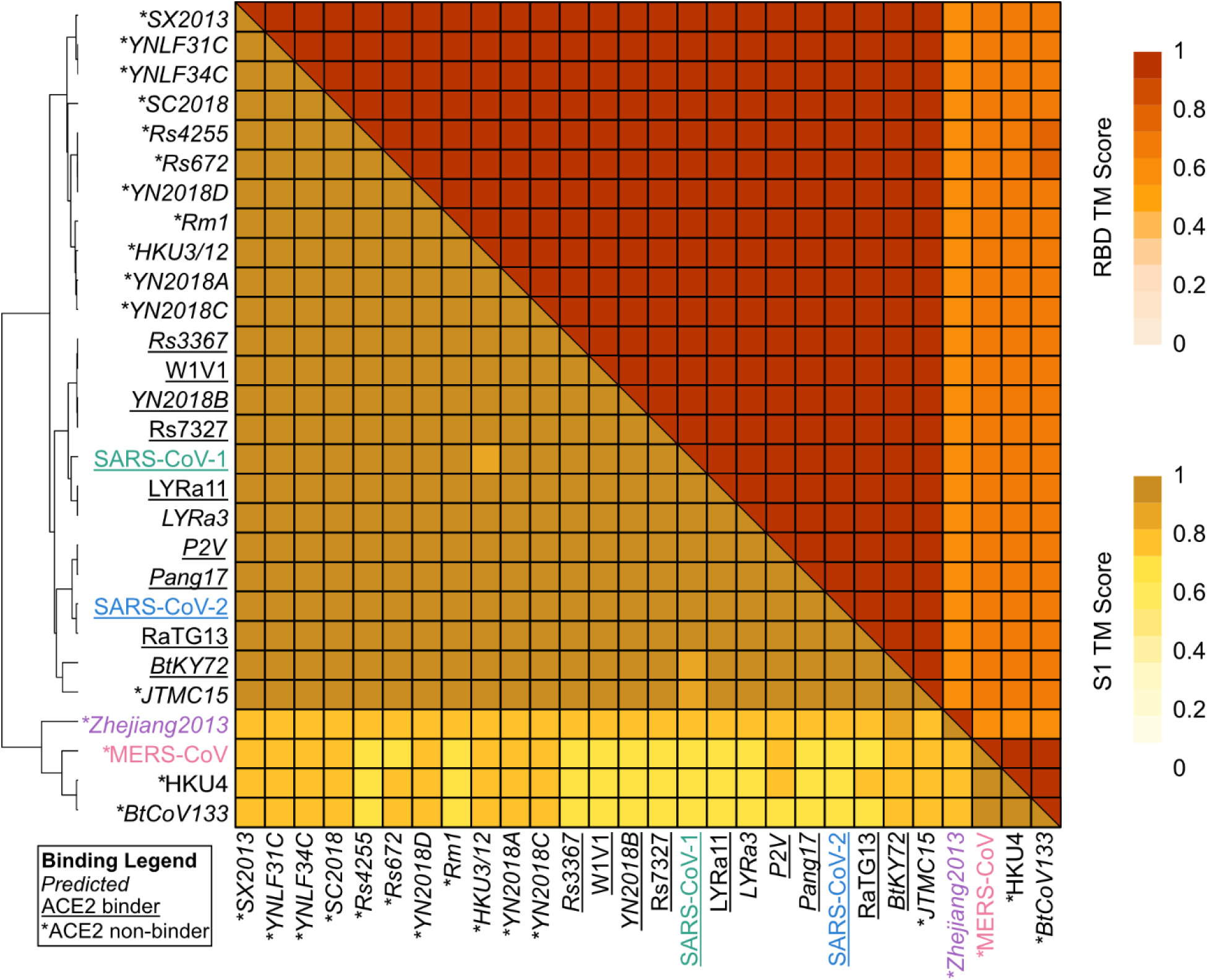
Structure analysis of emerging Spike S1 and RBD regions. The structure alignment heatmap of S1 and RBD shows the subgenera grouped together.

To achieve greater resolution of structural similarities and differences, we utilized the STRIDE web server, which implements a DSSP algorithm to quantify α-helices, β-strands, turns, and “other” (coils, 3₁₀-helices, and bridges). S1s were mostly comprised of β-strands (about 44%) (Figure S1A). RBDs exhibited a markedly higher proportion of intrinsically disordered regions, falling in the “other” category, with an average of about 39% (S1s averaged around 23%) (Figure S1B and Table S5). Nevertheless, there was no obvious correlation between S1 or RBD secondary structure content and known ACE2 binding.

### Predicting the S1-ACE2 interaction

We then predicted how each S1 might bind to human ACE2. Spike binds to its host receptor at the RBD located in the S1 domain, so we assumed that this function is maintained across other S1s. However, we reasoned that using S1 to model this protein-protein interaction may affect the total interaction energy and the algorithm’s prediction.

While there are many programs that can model protein-protein interactions, we used Multimer, ClusPro, and HADDOCK to predict interactions due to their ease of use, required inputs, and web-based interfaces. The results of each program gave varying degrees of interaction clarity when we analyzed the top-ranked model of each prediction (Figure S2). The lack of binding interface input for Multimer predictions meant that many of the predicted complexes were physically impossible, either due to the orientation of ACE2 or to how the S1 was folded with ACE2. As a structure prediction algorithm, Multimer does not support restraints. ClusPro makes restraints optional, and HADDOCK requires the information to simulate a protein-protein interaction. As with Multimer, unrestrained ClusPro predictions yielded physically impossible complexes. To guide the orientation of proteins in docking algorithms, we used restraints, or single residue pairings, to orient the RBD in S1 to the binding interface of ACE2 ^53^ (Table S6).

For ClusPro and HADDOCK, we used a combination of three amino acids in the receptor binding motif of each S1, paired with two amino acids at the top of ACE2, to orient the proteins into a physically feasible complex (Table S6). The ClusPro predictions with minimal restraints left more positional flexibility while keeping the proteins in more plausible configurations. The semi-flexible docking that HADDOCK affords resulted in more realistic complexes but can force interactions that do not normally occur (Figure S2).

To quantify protein binding, we calculated the total number of contacts between S1 and ACE2 for each platform (Figure 5 and Table S7). Contacts predict which amino acids may be interacting between molecules. Pandemic- and outbreak-causing viruses were used as controls to establish a binding threshold. Known ACE2 binders (SARS-CoV-1 and SARS-CoV-2) served as positive controls, whereas the known non-binder (MERS-CoV) served as a negative control. Contacts were calculated in ChimeraX, defining contacts as interatomic overlap of greater than - 0.4 Å. We categorized the viruses based on their predicted ability to bind ACE2 to assess whether we could infer binding from the number of contacts between the S1 and ACE2 ^31^. We categorized the hibecovirus as “N/A” since no predictions or experimental data have been reported for this virus regarding ACE2 binding.

**Figure 5.**
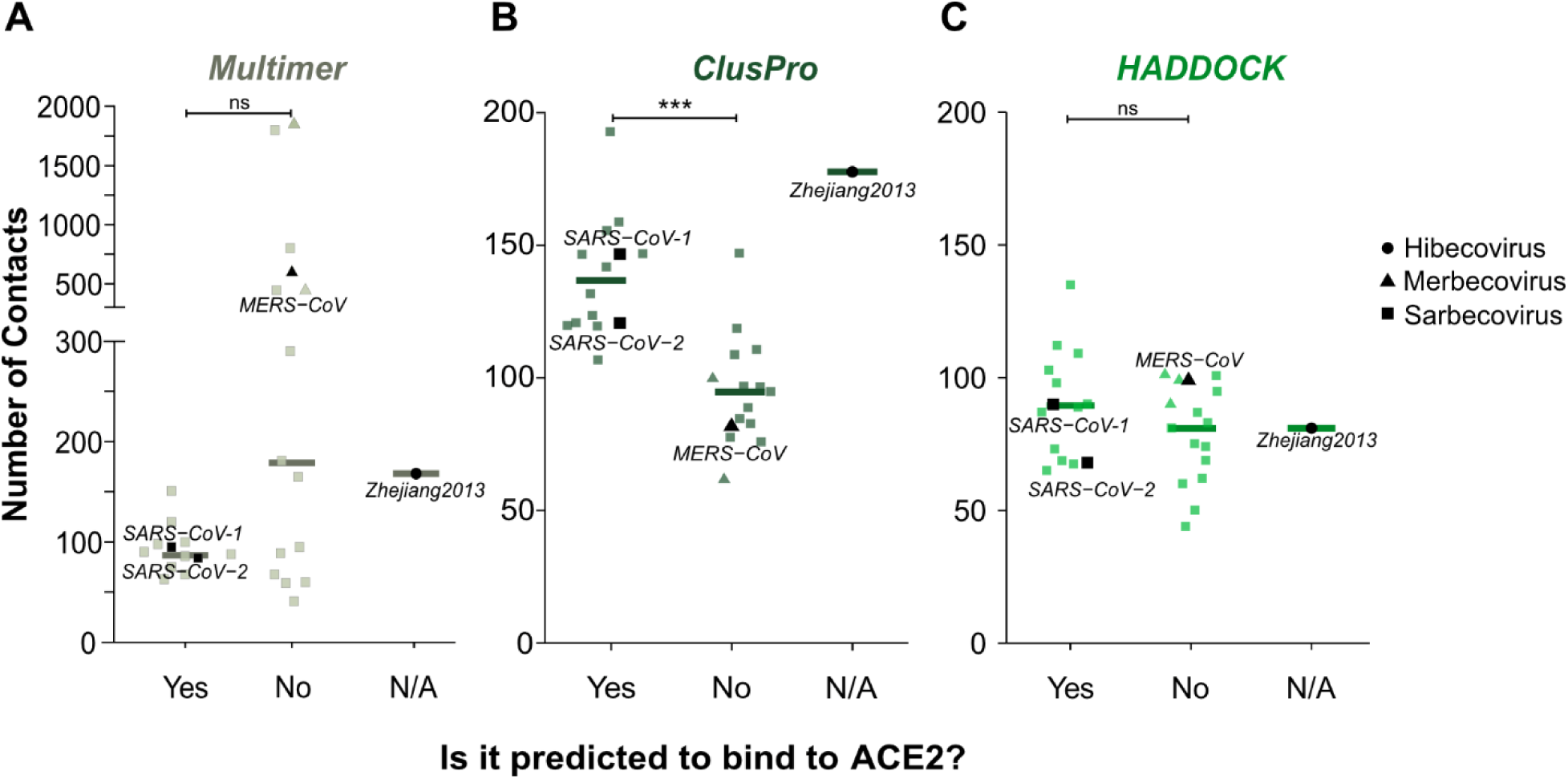
S1-ACE2 static interaction predictions show different binding behaviors. A-C) Number of contacts calculated between each S1 and ACE2 for A) Multimer, B) ClusPro, and C) HADDOCK. The bar indicates the median number of contacts for each subset. Significance was calculated using a Mann-Whitney U test, *** p < 0.001.

The protein-protein interaction prediction tools yielded distinct ranges of contacts for ACE2 binders and non-binders (Table S7). The number of contacts calculated for the Multimer predictions showed greater spread (Figure 5A), likely driven by physically unrealistic predictions for many viruses (Figure S2). For ClusPro, predicted/known ACE2 binders had significantly more contacts than predicted/known non-binders (Figure 5B). With HADDOCK, the number of contacts was not significantly different between these groups, possibly due to the greater number of restraints applied during docking, including non-binders (Figure 5C).

Since ClusPro best recapitulated the expected ACE2-binding behavior, we focused our additional analysis on predictions generated by this platform. We next analyzed the prediction for Zhejiang2013 more closely. This virus has not been studied, so its ability to bind ACE2 is unknown. Compared with known ACE2 binders and non-binders, the number of contacts between Zhejiang2013 and human ACE2 exceeded those of even SARS-CoV-1 and SARS-CoV-2 (Figure 5B and Table S7). This suggests Zhejiang2013 might be an ACE2 binder.

### Simulating the S1-ACE2 interaction using molecular dynamics

While ClusPro revealed consistent trends regarding ACE2 binders and non-binders, these predictions are limited by their static nature. To strengthen our confidence in the prediction that Zhejiang2013 S1 can bind human ACE2, we performed MD simulations. We also simulated SARS-CoV-1 and MERS-CoV, a known ACE2 binder and non-binder, respectively, as controls. We used coarse graining to enable extended simulation timescales ^54,55^. Additionally, to more directly compare and validate our static interaction predictions, we excluded the glycans known to decorate the Spike protein under normal conditions. While this oversimplifies the Spike-ACE2 protein-protein interaction dynamics, it provided a faster dynamic test of the interaction predictions ^56,57^.

The coarse-grained simulations recapitulated the overall behaviors observed by the contact analysis of the static predictions. SARS-CoV-1 S1 maintained continuous contact with ACE2 throughout the entire simulation (Figure S3). Interestingly, Zhejiang2013 S1 also remained bound to ACE2 for the full 1000 ns, supporting our hypothesis. MERS-CoV S1 dissociated from ACE2 in two out of three simulations. Analysis of root mean squared deviation (RMSD), which measures how much the complexes deviate from their initial conformations over time, showed that both SARS-CoV-1 and Zhejiang2013 had lower RMSDs than MERS-CoV, likely due to remaining in a complex like their original structure (Figure S4).

While coarse-grained models provided an initial indication of whether the S1 domains bind to human ACE2, they lack information about the electrostatic forces by reducing the atoms to beads. Therefore, we used all-atom steered MD (pulling) simulations to better understand the binding forces between Zhejiang2013 S1 and human ACE2 in comparison to SARS-CoV-1, a known ACE2 binder. Pulling simulations model the force required to pull two proteins apart, where a greater pull force indicates stronger binding ^58–60^. We did not simulate pulling of MERS-CoV S1 because it does not normally bind ACE2 and the two proteins did not stay bound in the coarse-graining simulations (Figure S3).

For these pulling simulations, we included glycans (Man5) on the proteins since glycans are known to influence pulling force required to dissociate SARS-CoV-2 RBD and ACE2, and regulate gate opening of the SARS-CoV-2 Spike trimer ^58,67^. Glycan positioning is well-known and experimentally resolved for pandemic-causing coronaviruses, but unknown for the emerging coronaviruses ^48,56,57,61,65,66^. Because Zhejiang2013 has not been studied experimentally, there is no experimental evidence available about its glycan positioning. We therefore predicted glycan positioning for both proteins using GLYCAM and NetNGlyc. We validated the positioning with the experimentally resolved glycosylation pattern of SARS-CoV-1 before continuing with Zhejiang2013 (Figure S5) ^58,61–64^.

We performed these simulations similarly to our previous work on SARS-CoV-2 by pulling the proteins at 1, 5, and 10 nm/ns pulling rates to assess robustness for this non-equilibrium process ^58^. We plotted the pull force as a function of pull distance between S1 and ACE2. The distances were calculated using the center of mass of each S1 and ACE2. We observed a higher peak pulling force for Zhejiang2013 compared to SARS-CoV-1 at all pulling rates (Figure 6 and S6). However, pulling force returned to 0 kJ/mol/nm for only the lowest pulling rate of 1 nm/ns. We therefore focused our analysis on this slower pulling rate since the simulation achieves force equilibrium. We observed that the complexes separated at around 8 nm and 10 nm for SARS-CoV-1 and Zhejiang2013, respectively, at this pulling rate (Figure 6). The curve for Zhejiang2013 S1 is also broader than SARS-CoV-1. The greater peak pulling force over a longer pulling distance indicates stronger binding of Zhejiang2013 S1 to ACE2 than SARS-CoV-1 at the binding interface.

**Figure 6.**
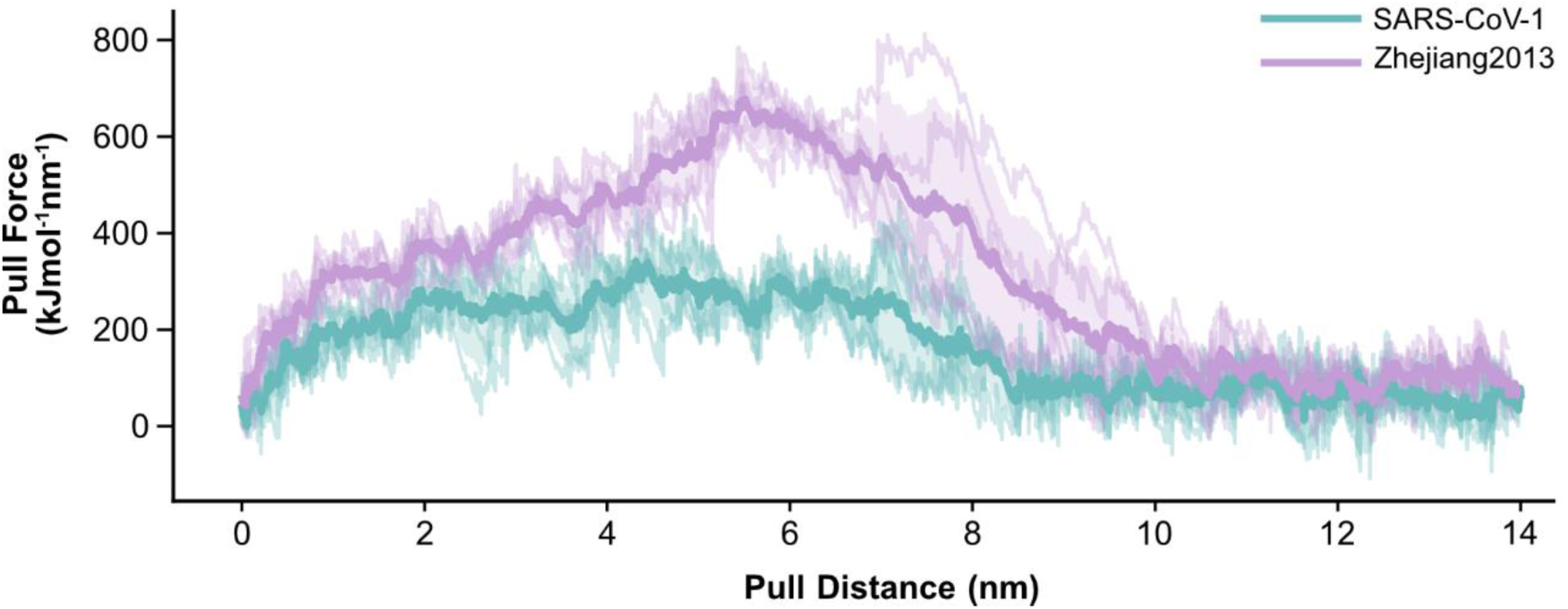
Pulling simulations reveal a stronger pulling force between Zhejiang2013 and human ACE2. Traces of pull force as a function of pull distance at a pulling rate of 1 nm/ns. We show five replicates per system and highlight the mean of the replicates. The shaded region shows the standard deviation.

## Discussion

Here, we screened various emerging betacoronaviruses to assess their potential to bind ACE2 using protein docking platforms and molecular dynamics simulations. We started by aligning the sequences to see how they might cluster and identify similar binding behaviors (Figure 2). We used pandemic- and outbreak-causing viruses as controls to set a binding threshold—SARS-CoV-1 and SARS-CoV-2 serving as positive controls for ACE2 binding, and MERS-CoV as a negative control. We then predicted 28 Spike S1 domains and aligned their structures to assess whether any conserved regions might mediate ACE2 binding (Figures 3 and 4). We predicted potential interactions using three approaches and quantified binding by counting the number of contacts between the proteins (Figure 5). ClusPro best recapitulated more contacts for known/predicted ACE2 binders versus non-binders. The number of contacts per S1-ACE2 interaction indicated that Zhejiang2013, a hibecovirus with unknown ACE2-binding ability, has more interactions with ACE2 than known ACE2 binders. To validate the docking predictions, we used MD simulations to examine the intermolecular forces at play and calculated the pulling force between S1 and ACE2 (Figure 6). Taken together, our results suggest that Zhejiang2013 has the potential to be an ACE2 binder, despite hibecoviruses not being described as ACE2 binders.

Focusing on the S1 domain offers a practical solution to overcome the difficulties of examining full-length Spike across different coronaviruses while maintaining a broader view that includes the RBD, especially for uninvestigated viruses. Studying the S1 region of Spike can open the possibility of identifying new binding modalities in cases where the RBD may be positioned slightly differently within the S1 structure ^32,46^. Finally, this approach, which encompasses a broader region of the viral protein than the known RBD, may also be important for viruses in families in which less is known about receptor binding. Thus, this more general approach retains flexibility for future pandemic virus risks.

We are not aware of any studies that test or predict the binding of a hibecovirus S1 to ACE2. Our computational modeling revealed unexpected similarities between hibecovirus Zhejiang2013 and SARS-CoV-1, a known ACE2 binder. The predicted S1 domain of Zhejiang2013 more closely resembled that of known ACE2 binders than non-binders. Both Zhejiang2013 and SARS-CoV-1 share a disordered region and an α-helix at the binding interface, which contrasts with the β-sheets in MERS-CoV at the same interface. These similarities indicated that Zhejiang2013 may retain structural elements necessary for ACE2 recognition, even if its genus is not usually associated with ACE2 binding. These results emphasize that small structural differences within the S1 domain can significantly influence receptor engagement.

This work also highlights the utility of easy-to-use protein interaction prediction platforms. Such platforms are valuable for screening emerging viral proteins and generating a reproducible screening pipeline. They are fast, computationally efficient, and require minimal prior information—a critical advantage for poorly characterized or uninvestigated proteins with limited experimental information to inform the interaction dynamics. Multimer only requires the protein sequence to predict a protein complex, but the lack of energy calculations to guide the prediction resulted in biologically improbable orientations. Conversely, HADDOCK provides a flexible docking approach for predicted structures but requires the positions of a known binding interface. Its predictions are more guided and thus more accurate for true interactions, but the requirement of a known binding motif makes it difficult to use for largely uninvestigated proteins. This could also “force” an interaction through a required or known binding interface when none should exist. ClusPro provided a “Goldilocks” solution by requiring only the predicted structure with some options for limited restraints. The capability of unrestrained docking, optimized by energy minimization, becomes a useful alternative for understudied proteins. ClusPro also provides many options to guide the docking if more information is available about the two proteins of interest. We found that using residue-based restraints afforded the predictions more flexibility and clustered a variety of possible orientations (Figure 5 and Table S6). This meant that we could discern statistically significant differences between predictions across the emerging S1s. Overall, protein interaction prediction platforms afford researchers quick methods to test protein binding. However, these tools generally treat proteins as rigid structures and ignore the temporal and structural dynamics of protein-protein interactions, which oversimplifies the model.

While ClusPro accurately predicted high/low contact interactions concordant with expected ACE2 binding, the initial protein docking results represent a first-pass computational screen at understanding a protein-protein interaction. The simplified structures plus the lack of temporal binding dynamics mean that docking captures only a snapshot of the interaction in a rigid position. For well-known proteins like SARS-CoV-1 Spike, we acknowledge that a glycan-free, static prediction oversimplifies the interaction with ACE2, and the S1-only model does not capture the full context of full-length Spike trimer. However, for emerging viral proteins aside from the controls, there is very little experimental data to inform a predictive simulation. Static docking levels the playing field by not requiring additional information and quickly calculating the most thermodynamically feasible complex between two proteins.

Our pulling simulations revealed that more pulling force is needed to disrupt the interaction between Zhejiang2013 S1 and ACE2 than SARS-CoV-1 S1 (Figure 6). This suggests that Zhejiang2013 may bind ACE2 strongly. In the context of pandemic screening, the ability to bind a known host cell receptor indicates a critical prerequisite for zoonotic spillover. In the simulations, Zhejiang2013 exhibited a binding profile much greater than SARS-CoV-1. This mirrors similar trends already measured with SARS-CoV-2 having a greater binding affinity to ACE2 than SARS-CoV-1 ^19,32,68–70^. Taken together, this suggests that Zhejiang2013 Spike may contain the essential features for human receptor usage.

Post-translational modifications (PTMs) play critical roles in protein folding, binding, and functions. Viral fusion proteins (i.e., class 1 viral fusion proteins like coronavirus Spike, HIV Env, and influenza HA) rely on PTMs like glycans for shielding important antigenic epitopes from host immune defenses, influencing target binding, increasing binding affinity, and stabilizing conformations ^57,58,71–75^. More specifically, for pandemic-causing coronaviruses, glycosylation at the RBD directly affects binding strength to the human receptor ^48,56–58,61,76^. However, structural data and residue positioning of PTMs can be inconsistent or missing, especially for less-studied proteins and viruses. To initially streamline our predictions, we did not include any glycans on the S1 domain since glycan positioning is not experimentally resolved for all the viruses we are screening. Additionally, we could not include PTMs on the S1 structures when we initially predicted them with AlphaFold. While this omission was necessary for the initial structure and interaction predictions and overall computational efficiency, we acknowledge that the glycan-free models do not fully reflect the complexity of binding. Our previous work showed while glycans are not required for RBD to bind ACE2, they can increase the binding range and affinity at the interface ^58^. As we continued with MD simulations, we predicted the positions of glycans across the protein to more accurately model the intermolecular binding forces (Figure S5). While these MD results provided a more accurate representation of binding forces, PTM positioning can be inconsistent. Additionally, the lack of experimental data on glycan positioning for Zhejiang2013 means that these results model a computational estimate rather than a confirmed structural feature. These findings position Zhejiang2013 as an important candidate for future structural characterization of its binding interface in the presence of relevant glycans. Additionally, the simulation data strongly suggest that Zhejiang2013 can bind ACE2 and attach to the host cell, but this does not guarantee successful viral entry or replication. This is simply the first step that must be satisfied for virus emergence and highlights the potential risk posed by Zhejiang2013.

Ultimately, this work focuses solely on computational methods for predicting binding between an emerging antigen and a human receptor. This pipeline demonstrates that computational screening of emerging coronaviruses can help to prioritize high-risk candidates for further experimental investigation. It combines different levels of computational rigor to screen and calculate potential binding to a human receptor. While this workflow simplifies interaction dynamics, it allows for screening of uninvestigated proteins and emerging viruses for which no prior information is available. By flagging Zhejiang2013 as a potential ACE2 binder, we can see how structural predictions can bridge the gap between screening a library of antigens for experimental validation and accelerating pandemic preparedness efforts with minimal resources.

## Conclusions

We predicted the structures of S1 proteins from 28 emerging coronaviruses screened them for pandemic potential by predicting their ability to bind to human ACE2. We quantified predicted protein-protein interactions using contact analysis, which highlighted the degree of imposed restrained determines whether a method can discriminate between binders and non-binders. Compared to the controls in the static predictions, we identified the potential binding of the hibecovirus Zhejiang2013, which was not previously known to bind ACE2. When modeled using steered MD simulations, Zhejiang2013 exhibited a higher peak pulling force and extended pulling time compared to SARS-CoV-1, consistent with similar or stronger ACE2 binding compared to known pandemic-causing coronaviruses. This approach highlights Zhejiang2013 as a candidate for future characterization of S1-ACE2 binding affinity.

## Acknowledgments

We would like to thank Dr. Adam Fishburn for his scientific feedback and support. AlphaFold predictions and US-align structure alignments were performed on the Franklin High Performance Computer servers at UC Davis. Funding to PSS and RF was provided by National Institute of Standards and Technology (NIST) through BioMade to UC Davis through the Rapid Assistance for Coronavirus Economic Response (RACER) project. SSG was awarded the Human and Animal Health research award by the UC Davis Designated Emphasis in Biotechnology program. GL was supported by the AMPAC Fine Chemicals Summer Research Internship and the Provost’s Undergraduate Fellowship. This work used Expanse at San Diego Supercomputer Center through allocation BIO250243 and BIO250277 from the Advanced Cyberinfrastructure Coordination Ecosystem: Services & Support (ACCESS) program, which is supported by U.S. National Science Foundation grants #2138259, #2138286, #2138307, #2137603, and #2138296.

## Methods

### Spike S1 Sequence and Structure Analysis

#### Sequence Alignment

The amino acid sequences of each S1 or RBD were aligned using ClustalOmega 1.2.4 web server using default settings ^77,78^. The resulting percent identity matrix was used to generate a heatmap and dendrogram using RStudio 4.3.3 using the R package *pheatmap* (v1.0.13) ^79^.

Dendrogram reflects hierarchal clustering of sequence identity patterns.

#### Protein Structure Predictions

Coronavirus Spike S1 domain structures as well as human ACE2 were predicted using AlphaFold2 (v2.3.2) on the UC Davis College of Biological Sciences High-Performance Computer (HPC) to increase the throughput of predictions ^50^. To prevent biased decisions and ensure reproducibility, we continued our analysis with the best-ranked model, a choice we consistently maintained throughout the computational analysis.

#### Structure Alignment

The PDB files from the AlphaFold2 structure predictions were uploaded to the US-align tool (version 20241201) ^51,52^ on the UC Davis College of Biological Sciences HPC to increase throughput of comparisons. The resulting TM-scores were used to generate a heatmap and dendrogram using R package *pheatmap*. Dendrogram reflects hierarchal clustering.

#### Structure Composition Analysis

To analyze differences in secondary structure composition among emerging coronaviruses, we used STRIDE ^80^, a web server for secondary structure assignments. STRIDE is based on the dictionary of secondary structure of proteins (DSSP) algorithm. The program generated an Excel file listing the proportions of α-helices, β-strands, turns, coils, 3₁₀-helices, and bridges, which we manually consolidated into four main classes: α-helices, β-strands, turns, and other (comprised of coils, 3₁₀-helices, and bridges). The same procedure was applied to the RBDs.

### Protein-Protein Interaction Predictions

#### Multimer

To predict an S1-ACE2 interaction, we input the amino acid sequences of each S1 and ACE2 into the AlphaFold Server, which is based on AlphaFold3, as separate molecules to model the protein-protein complex ^37,38^. We used the top-ranked model for further analysis.

#### ClusPro

ClusPro 2.0 docking was performed using .pdb files on a cpu server ^39–43,81^. S1 was the designated receptor labeled as chain A, and ACE2 was the designated ligand as chain L. For restrained docking, the file was generated using the restraint generator in ClusPro, which produces a .json file ^53^. We used the highest-ranked complex for further analysis.

#### ClusPro Docking Restraints

These restraints were based on structural and sequence alignments between each S1 and SARS-CoV-1 S1, using known key binding residues. Overall, the sarbecovirus S1 maintained the same residues between the structure and sequence alignment. However, the merbecoviruses and the hibecovirus had amino acid residues when aligned to SARS-CoV-1 S1 by its secondary structure or its protein sequence. For every program we used, we selected the highest-ranked model for subsequent analysis.

#### HADDOCK

HADDOCK v2.4 requires both protein structure files and interacting residues to guide the prediction of the protein-protein interaction ^44,45^. For each prediction, we uploaded each .pdb file and specified each molecule as a protein and specified the chain label—A for the S1 file and L for ACE2—and used default settings. We specified that the active residues of S1 were the protein’s RBD and the residues 26-48 and 318-358 on ACE2 to guide the correct orientation of the proteins.

#### Visualization and Contact Analysis

Protein structures were visualized and analyzed using UCSF ChimeraX 1.7.1 ^82–84^. Contacts were calculated as a Van der Waals overlap of greater than or equal to -0.4 Å between intramodel chains. Plots were visualized with R package *ggplot* (v3.5.1).

### Statistical Analysis

Statistical analysis was done using a two-way Mann-Whitney U test in RStudio 4.3.3.

### Coarse-Grained Simulations

We constructed coarse-grained complexes, where four atoms are represented by a single bead to preserve key properties while reducing the number of degrees of freedom, of the S1 subunit ACE2 for SARS-CoV-1, MERS-CoV, and Zhejiang2013. The transmembrane domain of ACE2 was excluded from all systems. Box preparation and solvation were performed using GROMACS 2025.1 ^85,86^. Minimization, equilibration, and production runs were performed using GROMACS 2020.4 on Expanse at the San Diego Supercomputer Center ^87–90^.

We adjusted the pH of each complex to 7.4 and used the MARTINI3 force field to coarse-grain the proteins ^91,92^. An elastic network was implemented to maintain the secondary and tertiary structures, with parameters set to -ef 700.0, -el 0.5, and -eu 0.9. The coarse-grained systems were then solvated in a cubic water box with at least 5 nm between the protein and the box edges, using 0.15 M NaCl. Energy minimizations were performed until the convergence criteria were met.

Energy minimization was then performed to remove steric clashes and relax the system. The minimization was carried out for 50,000 steps with an energy step size of 0.001 ps and a convergence criterion of 100.0 kJ/mol/nm (emtol), meaning that the maximum force on any atom was kept below 100.0 kJ/mol/nm. To verify the success of minimization, potential energy was plotted using Python, showing a steady decrease followed by a plateau.

Following minimization, equilibration at constant pressure of 1 atm using the Berendsen thermostat was performed using a molecular dynamics integrator with a timestep (*dt*) of 0.02 ps and *nsteps* = 2,500,000 ^93^. During this stage, the system was gradually brought to the target temperature of 310 K using the velocity rescale thermostat ^94^. The equilibration process was evaluated by monitoring the time evolution of density, kinetic energy, potential energy, pressure, and temperature.

For each S1–ACE2 complex, the production simulation was performed using a timestep (*dt*) of 0.02 ps and *nsteps* = 50,000,000, corresponding to a production time of 1 μs per simualtion. The simulations were performed under NPT conditions using the Parrinello-Rahman barostat for constant pressure at 1 atm and velocity-rescale thermostat for constant temperature at 310K ^94,95^. For each complex, three independent replicates were conducted. Trajectory data were recorded for subsequent analysis of the system’s structural and dynamical behavior. RMSD and RMSF were plotted for each simulation run.

### Steered All-Atom Molecular Dynamics

#### Glycosylation

We predicted the glycan positioning of SARS-CoV-1 and Zhejiang2013 with GLYCAM-Web, which uses GLYCAM Molecular Modeling Library v1.5.1., and NetNGlyc-1.0 web interfaces ^62,64^. Both programs predicted the same positions, though NetNGlyc predicted more glycan positions than GLYCAM on Zhejiang2013. We continued with GLYCAM predictions for a more conservative prediction. The GLYCAM output for SARS-CoV-1 was compared to experimentally resolved data for validation ^96–98^. and corroborated those positions with experimental data. We added Man5 glycans to the S1–ACE2 complex of each viral variant, using the GLYCAM web interface. The ClusPro-docked S1–ACE2 complex was used as input for glycan attachment. ACPYPE (v2023.10.27) was used to convert the GLYCAM output files into GROMACS-compatible formats ^99,100^.

#### Pulling Simulations

The S1–ACE2 complex was simulated with the Amber ff99sb forcefield and placed in a simulation box and solvated using the SPC/E water model and 0.15 M NaCl, while maintaining a minimum distance of 3 nm between the solute and the box edges, resulting in a box dimension of 20 x 20 x 40 nm ^101,102^. Box solvation was performed in GROMACS 2025.1. After the box was generated, the complexes were rotated so that the pulling direction was along the x-axis. Then we performed a steep minimization step of 50,000 steps with an energy step size of 0.01 ps and a convergence criterion of 1000.0 kJ/mol/nm (emtol). This was followed by an NPT equilibration step using a molecular dynamics integrator with a timestep (dt) of 2 fs and nsteps = 50,000. Minimization, equilibration, and production were performed in GROMACS 2020.4 on Expanse.

Pulling simulations were performed under NPT conditions using 2 fs timesteps, a Nose-Hoover thermostat at 310K, and a Parrinello-Rahman barostat at 1 atm ^95,103^. In all runs, ACE2 was restrained while S1 was pulled with a spring constant of 1000 kJ/mol/nm^2^. A total of 30 pulling simulations were performed at three different pulling rates (1, 5, and 10 nm/ns) over 14 nm. Simulations were repeated five times and averaged for each complex.

