## Supplemental Information for "Discriminating betacoronavirus receptor usage across subgenera using protein structure prediction and molecular dynamics"

**Supporting Information**

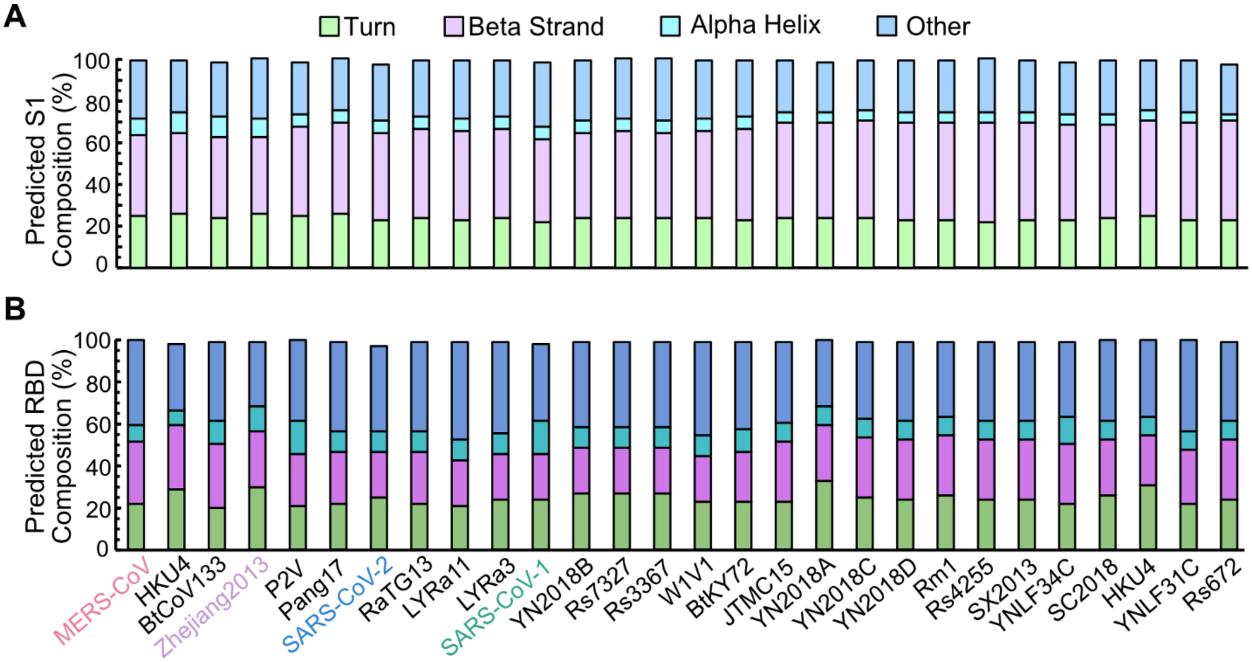


**Figure S1. Secondary structure composition using STRIDE.** Percentage of secondary structure predicted components of A) S1 and B) RBD.

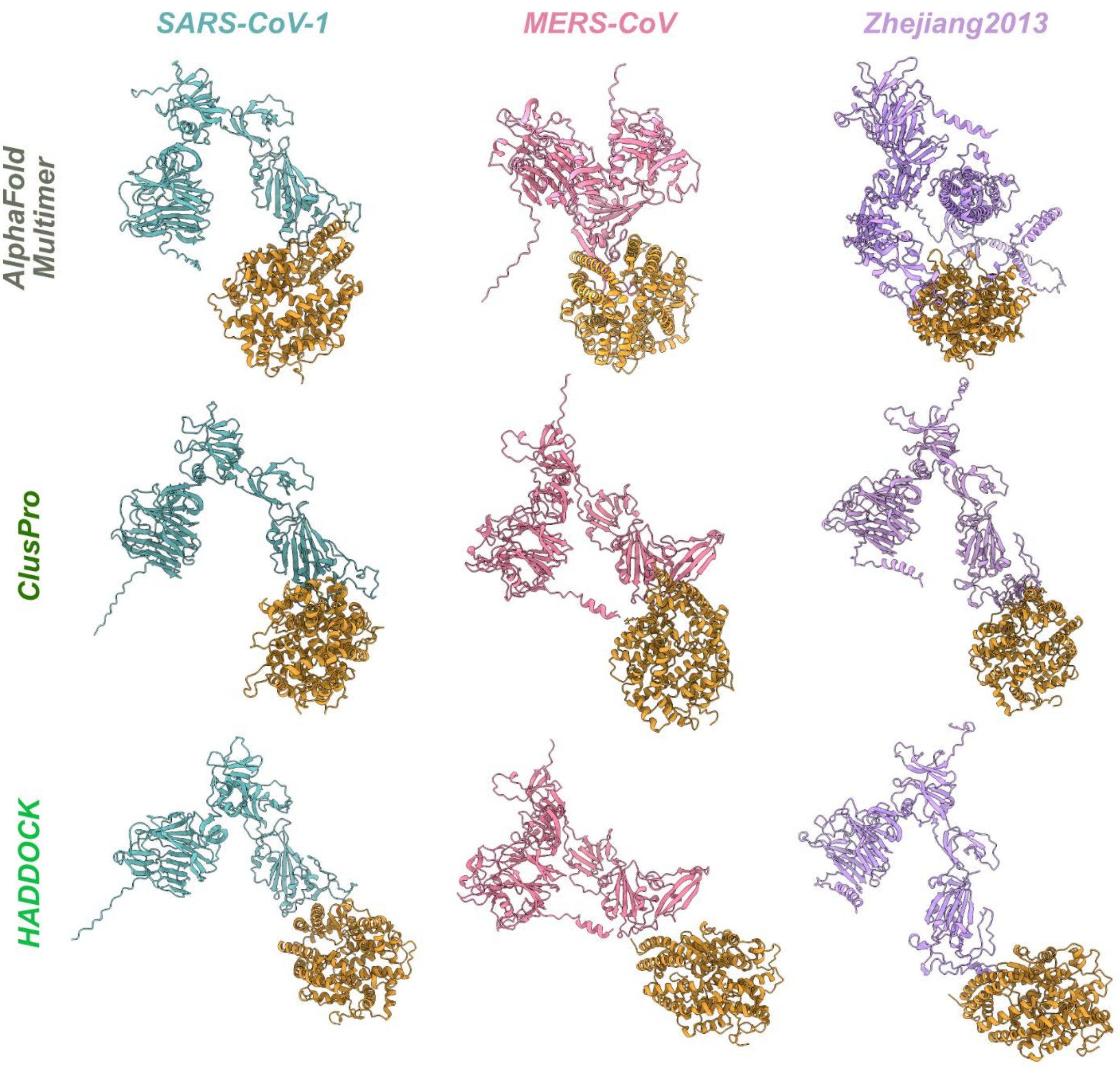


**Figure S2. S1-ACE2 static predictions of AlphaFold Multimer, ClusPro, and HADDOCK.**

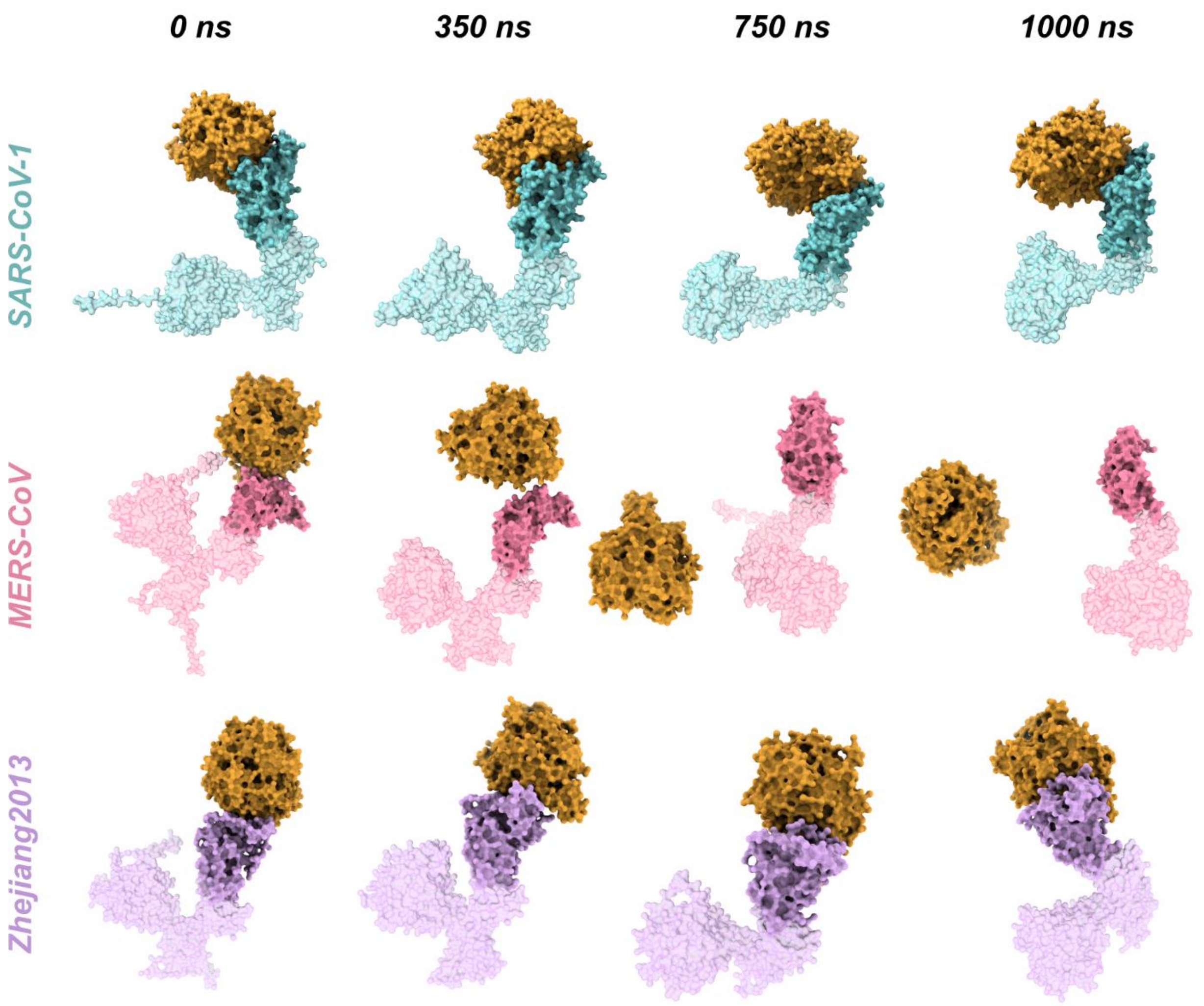


**Figure S3. Coarse-graining montage across a 1000 ns simulation of SARS-CoV-1, MERS-CoV, and Zhejiang2013 S1 binding to human ACE2.** Representative frames of a single trajectory are shown for each virus. SARS-CoV-1 (teal) and Zhejiang2013 (purple) remain attached to ACE2 (gold) at the RBD, whereas MERS-CoV (pink) is detached from ACE2. The RBD is emphasized as more opaque than the lower transparency S1.

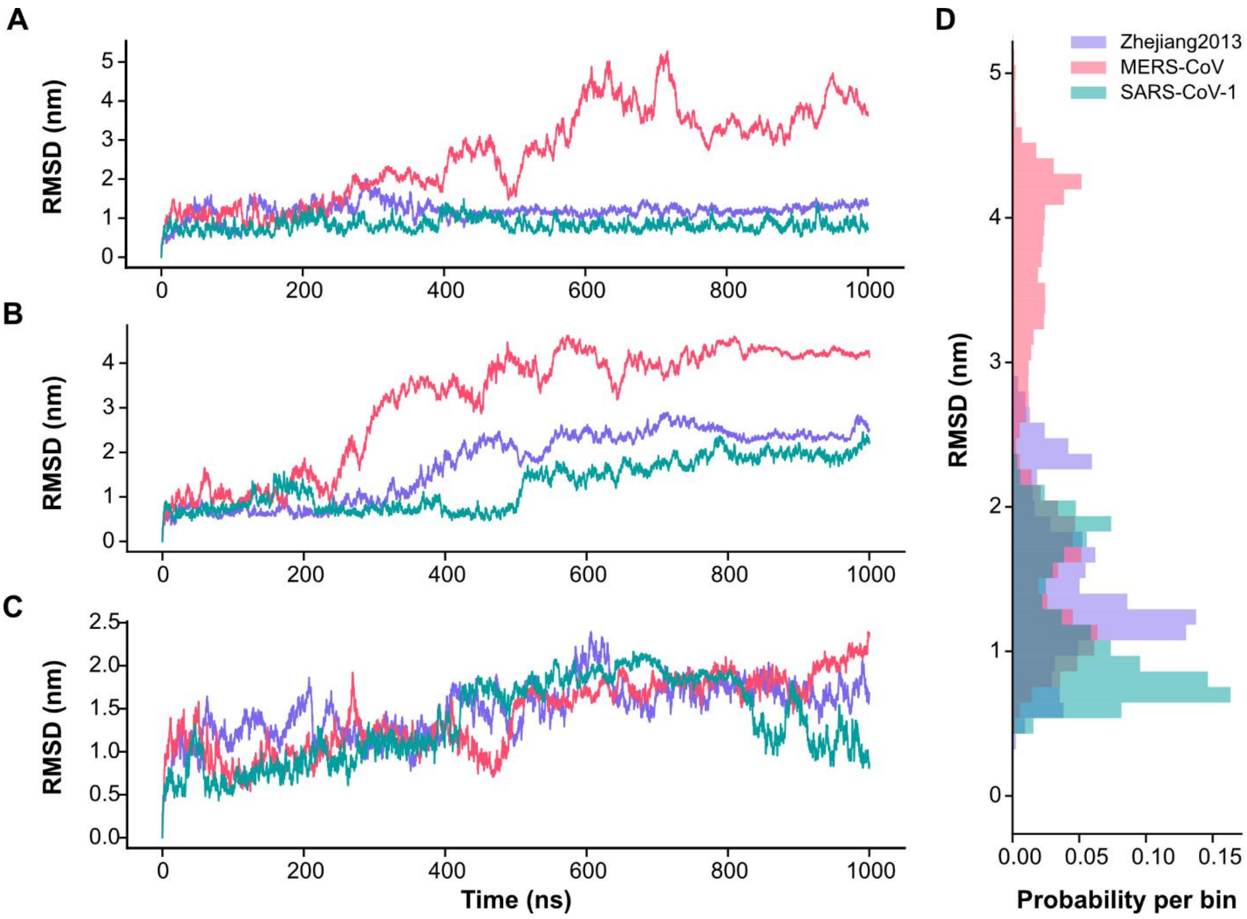


**Figure S4. Root mean squared deviation (RMSD) of coarse-grained simulations**. A-C) RMSD plotted as a function of time in nanoseconds across three replicates for each virus: SARS-CoV-1 (teal), MERS-CoV (pink), Zhejiang2013 (purple). Two simulations (A and B) resulted in MERS-CoV detachment from ACE2 and a higher RMSD over time. D) RMSD plotted against probability highlights the consistency or variability of protein behavior across all three replicates for each virus.

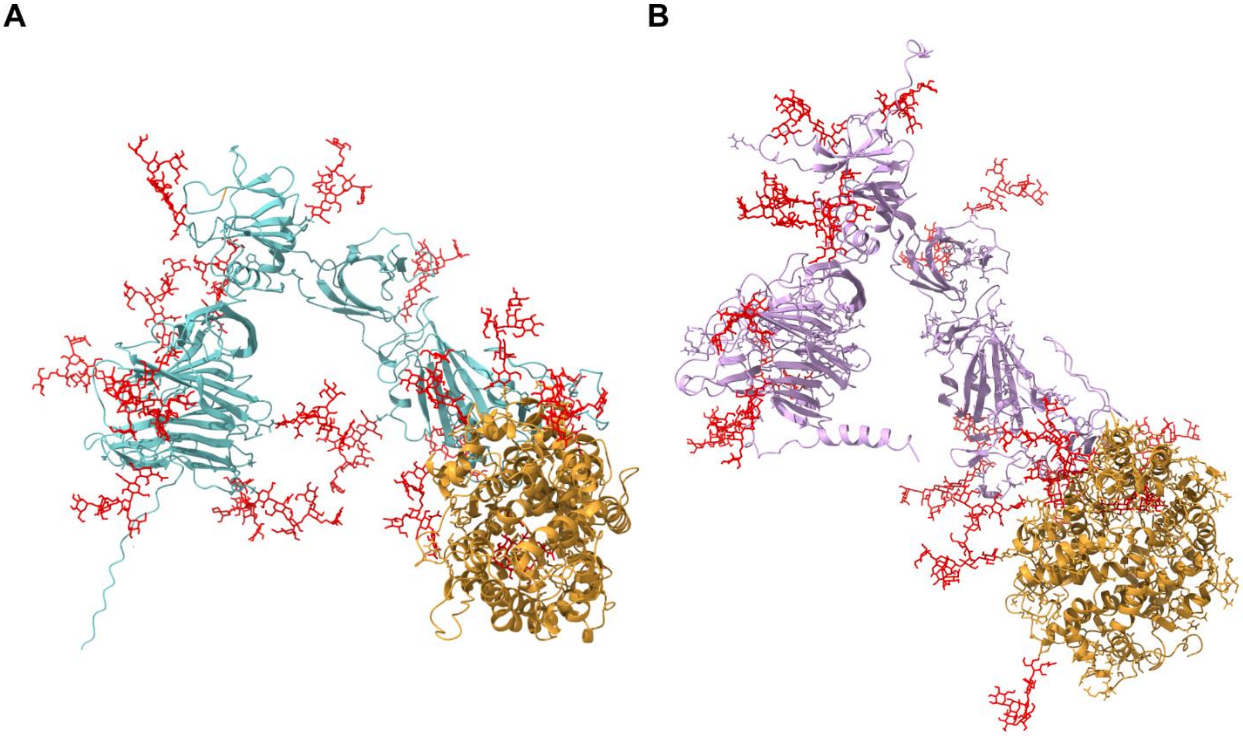


**Figure S5. Predicted glycosylated structures of S1 bound to ACE2.** For both A) SARS-CoV-1 and B) Zhejiang2013, the Man5 glycan (red) positions were added using GLYCAM.

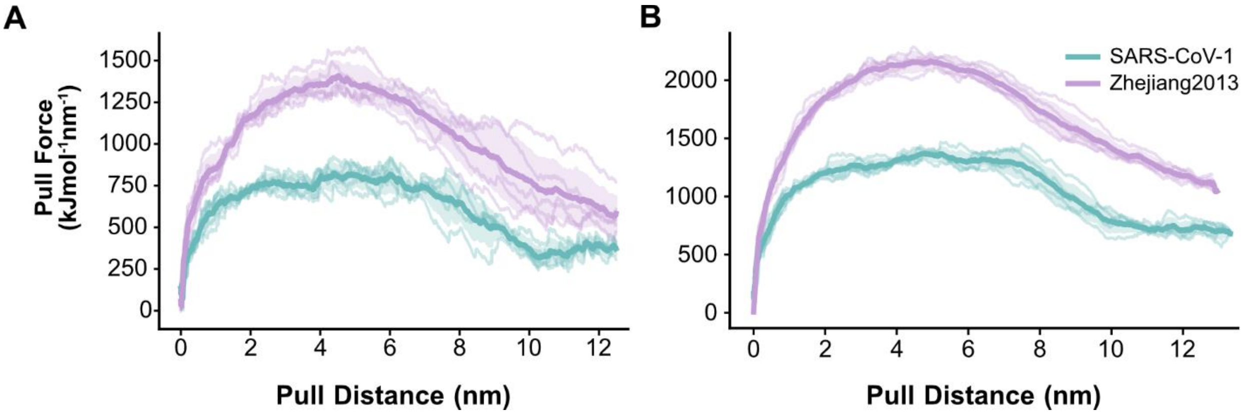


**Figure S6. Traces of all-atom steered MD simulations.** Plots compare force–distance profiles for Zhejiang2013 and SARS-CoV-1 at a pulling rate of A) 5 nm/ns and B) 10 nm/ns. We plot five replicates per system and highlight the mean of the replicates and shade the standard deviation region.
